# Cultural specialization as a double-edged sword: division into specialized guilds might promote cultural complexity at the cost of higher susceptibility to cultural loss

**DOI:** 10.1101/2022.04.22.489202

**Authors:** Yotam Ben-Oren, Oren Kolodny, Nicole Creanza

**Affiliations:** Department of Ecology, Evolution, and Behavior, Silberman Institute for Life Science, The Hebrew University of Jerusalem; Department of Biological Sciences, Vanderbilt University

**Author notes:** These authors contributed equally (Oren Kolodny, Nicole Creanza).

## Abstract

The transition to specialization of knowledge within populations could have facilitated the accumulation of cultural complexity in humans. Specialization allows populations to increase their cultural repertoire without requiring that members of that population increase their individual capacity to accumulate knowledge. However, specialization also means that domain-specific knowledge can be concentrated in small subsets of the population, making it more susceptible to loss. Here we use a model of cultural evolution to demonstrate that specialized populations can be more sensitive to stochastic loss of knowledge than populations without subdivision of knowledge, and that demographic and environmental changes have an amplified effect on populations with knowledge specialization. Finally, we suggest that specialization can be a double-edged sword; specialized populations may have an advantage in accumulating cultural traits but may also be less likely to expand and establish themselves successfully in new demes due to the increased cultural loss that they experience during the population bottlenecks that often characterize such expansions.

## Introduction

Throughout history, human culture has tended to accumulate complexity over time [1–3], and many studies have sought to measure or predict cultural complexity in different populations [4–11]. Multiple factors have been suggested as predictors of cultural repertoire size including population size, population mobility, the resources used by the population, and the rate of fluctuation of the environment [7,12,13]. One major transition in human cultural evolution that could have facilitated rapid cultural accumulation is specialization, when subsets of a population become proficient at making certain tools and accumulate relevant knowledge that is not shared by the rest of the population [14,15]. For example, the progression from non-specialized hunting-gathering to hunting with specialized tools has been proposed as a hallmark of the Middle Paleolithic to Upper Paleolithic transition, which has been characterized by a rapid diversification of cultural artifacts [16–19]; populations with a diverse set of specialized tools could putatively have an advantage if individuals themselves specialize in the production of these tools [20–23].

Imagine a population in which individuals produce cultural innovations (“tools”) at a certain rate. Given that the population’s repertoire is the number of unique tools across all individual toolkits, the smaller the overlap between those toolkits, the larger the population’s cultural repertoire will be (Figure 1). Thus, a *non-specialized* population, in which all individuals have similar toolkits, may be expected to have a smaller total repertoire than, say, a population divided into ten guilds that each have a different cultural specialization.

**Figure 1.**
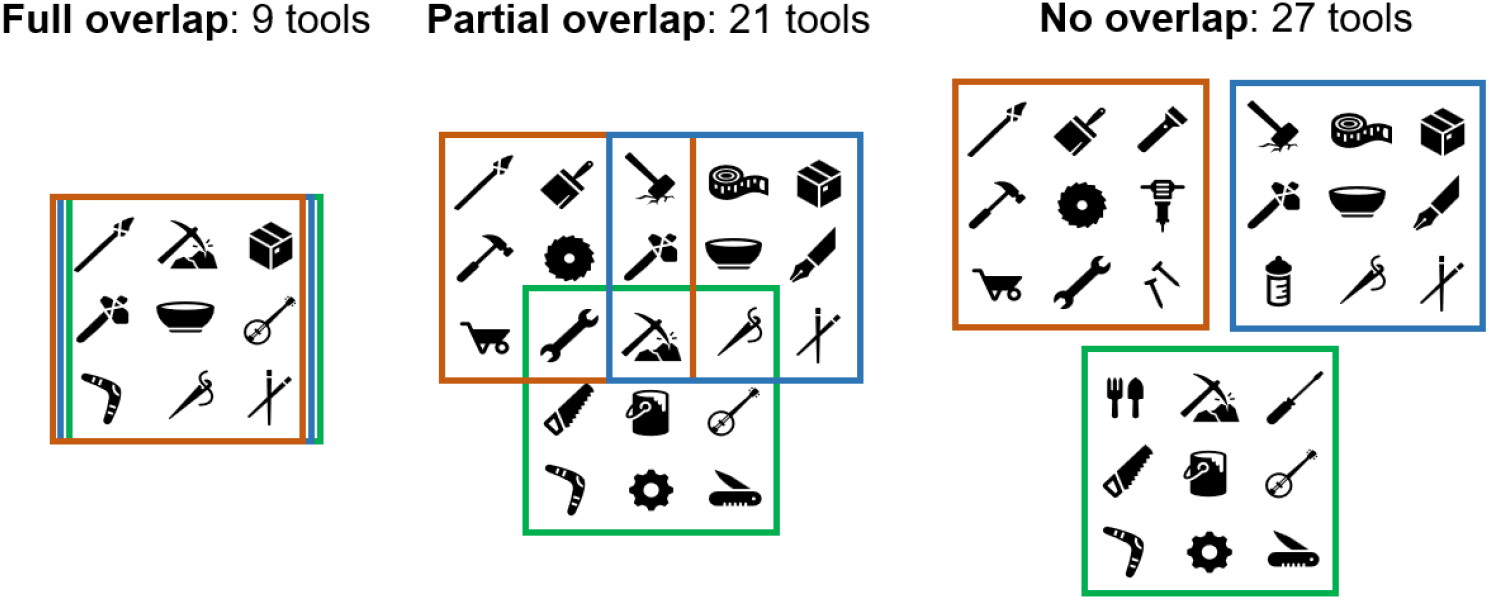
The smaller the overlap between individual toolkits (in this case, three individuals each with a toolkit of nine tools), the bigger the population’s overall cultural repertoire can be. At the same time, smaller overlap also means fewer people know each tool, which can increase its chances to be lost.

Thus, if small groups of individuals specialize in various domain-specific subsets of knowledge, a population can dramatically increase its overall cultural repertoire without requiring that members of that population increase their individual capacity for knowledge. However, specialization might be a double-edged sword in cultural accumulation, since cultural knowledge that is known by only a small subset of a population could be more likely to be lost, particularly during periods of demographic change [15]. In this paper, we present a computational model to test the assumptions underlying this key question: whether a subdivision of knowledge can lead to a richer cultural repertoire, and whether specialized populations can be more vulnerable to loss of knowledge.

Underlying these ideas is the assumption that larger populations can accumulate larger cultural repertoires, which has been the subject of recent debate [13,24–29]. Different approaches have been used to examine this assumption: some empirical analyses of the archaeological record and of existing populations have shown a correlation between a population’s size and the size of their cultural repertoire, but others did not find such a correlation [9,29–34]. Experimental studies of human innovations have shown an association between population size and innovation and also found an increase in innovations in partially connected social networks, in which individuals are connected to a subset of others [35–37]. Finally, several computational models, using different approaches and for different considerations, predict a positive relation between population size and cultural repertoire size [27,38,39]. None of these approaches has led to a general agreement. Notably, it has been suggested that a population size–repertoire size relationship is more often observed in food-producing populations, which may have more culturally mediated buffers against environmental fluctuations than hunter-gatherer populations [12,13].

The degree of cultural specialization of a population might thus be an important factor to consider in this debate, since a more specialized population may have a different level of cultural complexity than a non-specialized population at the same population size. In other words, considering populations with different degrees of cultural specialization in the same analysis could obscure the relationship between cultural repertoire size and population size. Here, using a model of cultural evolution, we examine the relationship between cultural specialization, cultural repertoire size, and sensitivity of the repertoire to stochastic losses and to demographic changes.

To conduct a preliminary assessment of the empirical correlation between population size and the degree of craft specialization, we analyzed data from the Ethnographic Atlas [40], digitized in D-PLACE [41]. There were eight cultural specializations listed in the database (metal working, leather working, house construction, pottery making, boat building, animal husbandry, hunting and fishing), and we assessed that each of these was a craft specialization in a population, coded in the database as “Craft specialization, i.e., the activity is largely performed by a small minority of adult males or females who possess specialized skills”, suggesting that related tools are known to only a subset of the population. We tallied the number of such craft specializations in each population for which population size estimates were available in the Ethnographic Atlas (953 out of the 1291 populations) and assessed the correlation between these two variables with a Spearman’s rank correlation.

## Methods

We propose a model of cultural evolution in which cultural traits, hereafter referred to as “tools,” occur in specialized and in non-specialized populations. The degree of population specialization is defined as the number of guilds into which it is divided. Non-specialized populations, hence, are considered as consisting of one guild. The cultural dynamics within each guild are completely independent from other guilds, meaning there are no shared tools between them.

Our model is roughly based on the modeling framework of [42] for independent cultural innovations, termed main axis tools in the original model, with adjustments for the division into guilds. In the model presented here, new tools are invented at a given rate per individual per time-unit, which was set at 0.001, similarly to the original model. This means the number of invented tools per time-unit is linearly dependent on population size (regardless of division to guilds). Realistically, the relation can exceed linear (as demonstrated in [42]) because existing tools can be used to invent new tools, but for simplicity, our version of the model assumes the inventions of tools are independent from one another.

Invented tools can be lost in two ways. First, newly invented tools can be lost before being established in the population. Each new tool is associated with a positive selection coefficient *s*, which is drawn from an exponential distribution with parameter β = 0.01. We do not explicitly simulate the cultural transmission process that would result in this type of loss; instead, as in [42], we use an approximation that a newly invented tool is stochastically lost with probability 1−*s* [43]. Thus, with probability *s*, a newly invented tool is established and is immediately considered to have reached its equilibrium frequency in the population.

In addition, after a tool has been established in the population, it can still be lost due to drift. Kolodny et al. did not attend to the issue of specialization [42,44,45], and therefore did not consider each individual’s potential capacity to accumulate cultural traits or the level of overlap between individual repertoires [11,46]. Hence, in the original model, the probability of loss of an individual tool was in negative linear relation to the population size.

In our model, we consider that the repertoire size may also be limited by each individual’s capacity to accumulate cultural traits. We thus assume that the probability of loss of a tool is also exponentially dependent on the existing repertoire size. That is, the bigger the cultural repertoire, the harder it becomes to remember all tools and transmit them to the next generation. The loss function we used is 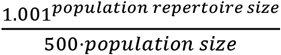.

We also examined alternative approaches to simulate the relation between the existing repertoire and the probability of loss, which yielded similar results. One such example (demonstrated in Figure S1) was to assume that probability of loss of tools is independent of repertoire size (as in [42]), but that there is a maximal repertoire that a guild can achieve, which is derived from the individual capacity for knowledge. This follows from the assumption that within a guild, most individuals know most tools, so the total repertoire of a guild cannot greatly exceed what one person can learn individually.

In our simulations, the cultural repertoire size of a population plateaus, reaching a steady state at which the number of tools fluctuates stochastically around a stable value. The repertoire size of a population at this steady state, termed here as average cultural repertoire size, was estimated by averaging the observed repertoire of a population between time steps 200,000 (at which it has already plateaued) and 500,000. In cases when we measured the average cultural repertoire at different time points in a simulation, such as before and after a bottleneck, we similarly averaged over 300,000 time steps after the repertoire size had plateaued.

First, we examine the relationship between population size and repertoire size in non-specialized populations; then, we examine the average cultural repertoire sizes achieved by different numbers of guilds for different population sizes. Increasing the level of specialization can potentially allow a population to maintain a larger cultural repertoire, but subdividing into too many guilds might lead to increased cultural losses as the number of individuals in each guild decreases. The number of guilds that leads to the largest average cultural repertoire size of a population is termed the repertoire-maximizing number of guilds.

Assuming a guild that loses a high percentage of its repertoire within a short time frame may collapse, highly specialized populations composed of many small guilds may not be sustainable. Thus, we compare repertoire-size stochasticity for guilds of different sizes.

Finally, we demonstrate how demographic and environmental changes might affect repertoire size differently in non-specialized and specialized populations. In all analyses in this section, we start by allowing both populations to reach a steady-state repertoire size. In the first analysis, we then let population size fluctuate: with a default probability of 0.00001 per time step, the population can increase or decrease by 200 individuals, with the total population size constrained to between 600 and 1000 individuals. In the second analysis, we impose a fluctuating environment. We define a probability of 0.0001 for shift between two environments, assuming that 90% of tools invented in a given environment are only useful in that environment, which results in a 10-fold increase in their probability of loss in the alternative environment.

## Results

### The relationship between population size and repertoire at equilibrium

The relationship between population size and average cultural repertoire size in a non-specialized population (consisting of one guild) is sigmoid-like in our model (see Figure 2). For small population sizes, the relationship has an increasing slope, e.g. doubling the population size from 10 to 20 quadrupled the number of tools from 5 to 20, and at larger population sizes has a decreasing slope, e.g. doubling population size from 10,000 to 20,000 increases the number of tools by a factor of only 1.18, growing from 6630 to 7850). Given this relationship, we can predict when the average cultural repertoire of a specialized population in our model is expected to exceed that of a non-specialized population. Whenever doubling the population size from N to 2N less than doubles the average repertoire size (which starts happening around N = 170 individuals) it means a non-specialized population of 2N individuals will have a higher average repertoire if it divides into 2 guilds.

**Figure 2.**
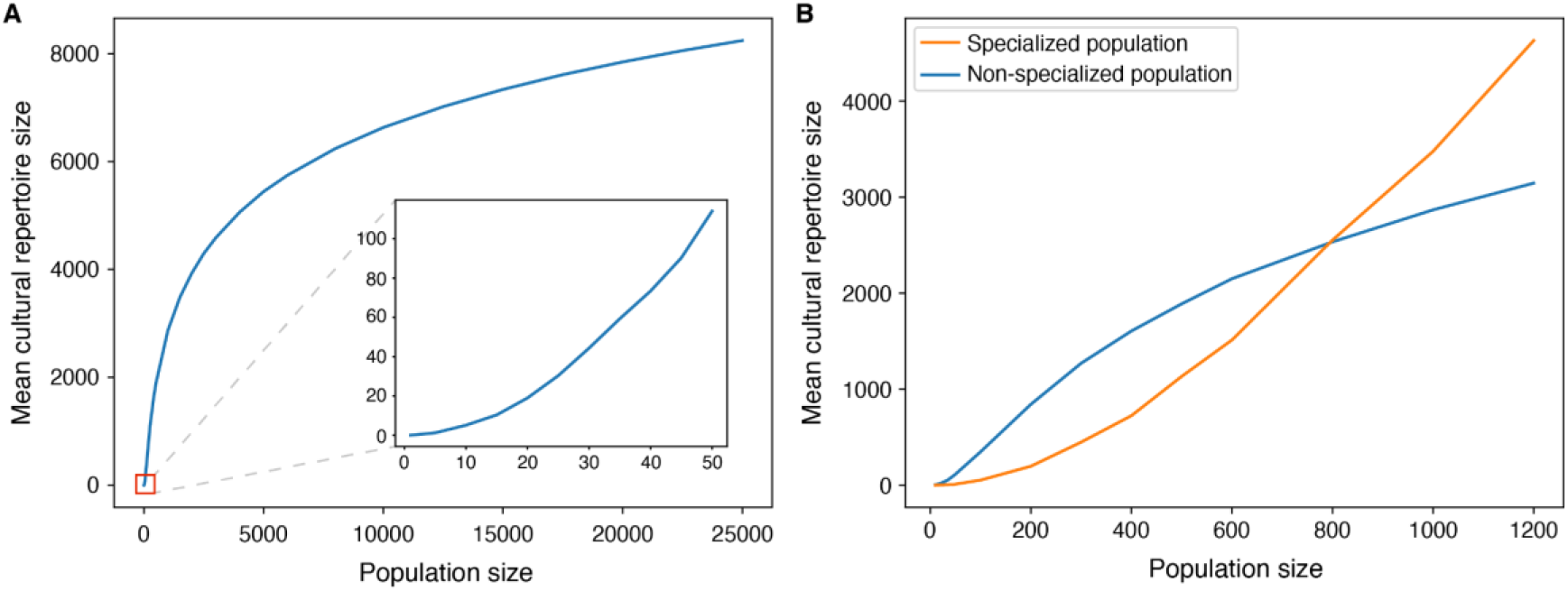
The relationship between population size and average cultural repertoire size in our model. **(A)** In small non-specialized populations (see the zoomed panel), the relationship has an increasing slope, which decays as population sizes increase. **(B)** At small population sizes, non-specialized populations can reach higher average repertoire sizes, while in larger populations, specialized populations (in this case, divided to 10 guilds) surpass them. Here, specialized populations with 10 guilds have larger cultural repertoires than non-specialized populations only in populations with more than ∼800 individuals. For specialized populations with fewer guilds, this threshold population size would be smaller.

Thus, we examine the number of guilds that will maximize the average repertoire size for populations of different sizes (Figure 3). We show that at smaller numbers of individuals, non-specialized populations achieve higher repertoire sizes than specialized ones, but as populations get larger, the repertoire-maximizing number of guilds increases.

**Figure 3.**
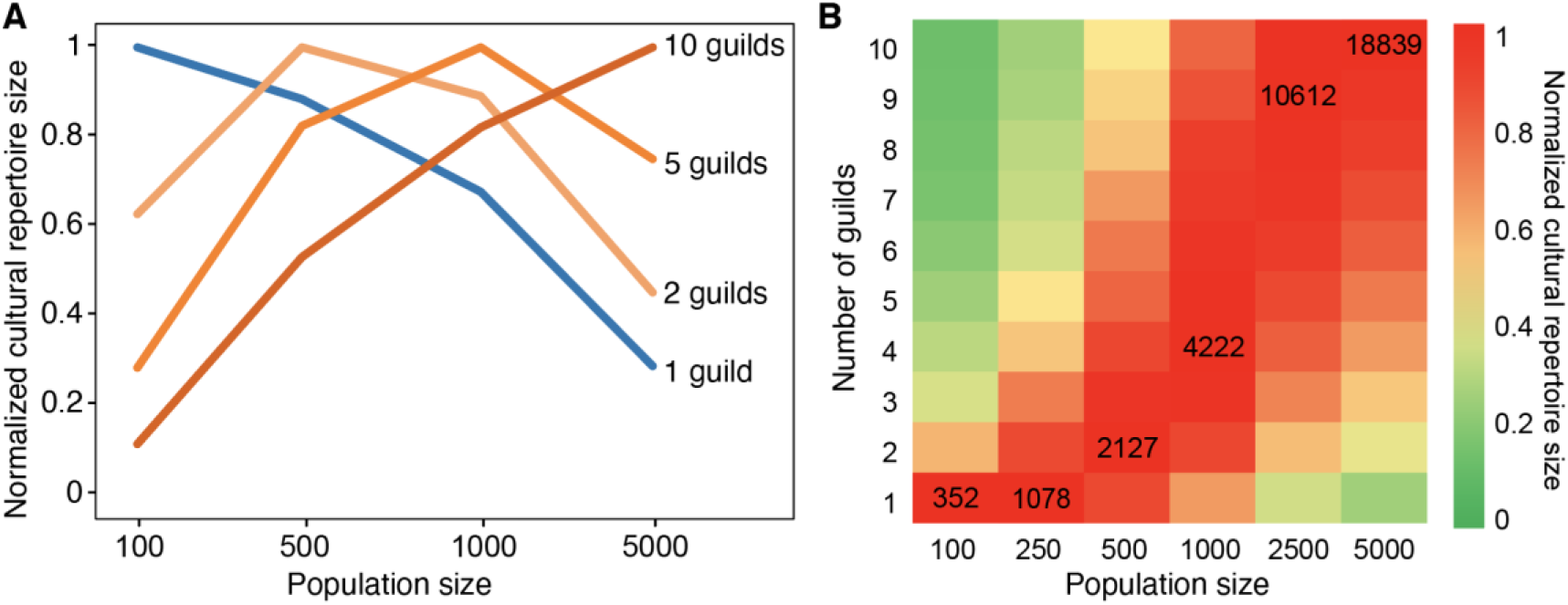
**(A)** A comparison between the relative repertoire sizes of populations that are either non-specialized (1 guild) or specialized (2, 5, or 10 guilds), for different population sizes. For each population size (100, 500, 1000, or 5000), each population’s average repertoire size was divided by the highest average repertoire size achieved for that population size. **(B)** A heat map showing the repertoire-maximizing number of guilds for different population sizes. The size of the largest cultural repertoire is indicated in each population size column; as in panel A, all cultural repertoire sizes in a given column are divided by this number such that the largest repertoire size is set to 1. For smaller populations in this simulation (100 and 250), non-specialized populations have larger repertoires than specialized populations with the same number of individuals, but for larger populations (500 and above) specialized populations have larger repertoires. Note that in both panels of this figure, for comparability, the repertoire sizes of each population were normalized by the maximal number of tools found in a population of that size. Thus, the value shown for the maximal number of tools, for each population size, is 1. The figure allows comparison of the tool repertoire achieved under scenarios of splitting to different numbers of guilds for each given population size, but not comparison of absolute repertoires of populations of different sizes.

Next, we examine the relationship between the size of a guild and the stochasticity in its repertoire at equilibrium (Figure 4). We show that the bigger the population, the smaller the relative stochasticity. We run simulations for populations sized 10, 100 and 1000 for 0.5 million time steps and measure the difference between the maximal and minimal population size achieved at equilibrium, i.e. the highest and lowest number of tools observed during the steady state between time steps 200,000 and 500,000). We find that for a guild of 10 individuals, the maximal difference in repertoire size was 100% (from a maximum of 12 tools to a minimum of 0 tools); for a guild of 100 individuals it was 22.5% (from a maximum of 395 tools to a minimum of 306 tools) and for a guild of 1000 individuals it was 6.7% (from a maximum of 2964 tools to a minimum of 2766 tools).

**Figure 4.**
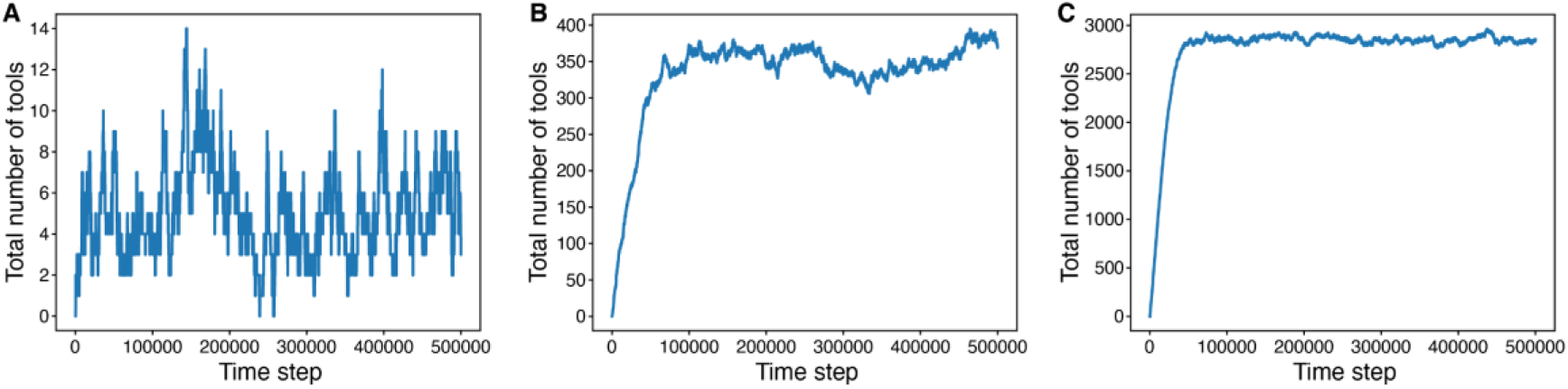
Relative cultural stochasticity is increased in small guilds. Panel **(A)** shows repertoire fluctuations in a guild sized 10; **(B)** in a guild sized 100 and **(C)** in a guild sized 1000.

### The effects of demographic and environmental changes on non-specialized vs. specialized populations

We compared the effect of fluctuations in population size on a non-specialized population and a specialized (10-guild) population (Figure 5). To make the results straightforward to compare, we chose to begin from a population size of 800 individuals, for which the average repertoire sizes will be similar between both populations (i.e. where the lines in Figure 2B intersect). For this population size, both populations plateaued around ∼2500 tools. After the repertoires reached equilibrium, we then allowed the population size to vary between 600 and 1000 individuals. We found that the corresponding variance in repertoire size was larger in the specialized population (varying between 1520 and 3580 tools, while the non-specialized population varies between 2140 and 2850). Although we chose an initial population size of 800 to examine the degree of cultural repertoire fluctuations when a non-specialized and specialized population each had a similar repertoire size, different population sizes showed similar results: specialized populations had larger relative fluctuations in cultural repertoire size than non-specialized populations experiencing the same changes in population size, regardless of the mean repertoire size of the population (Figure S2).

**Figure 5.**
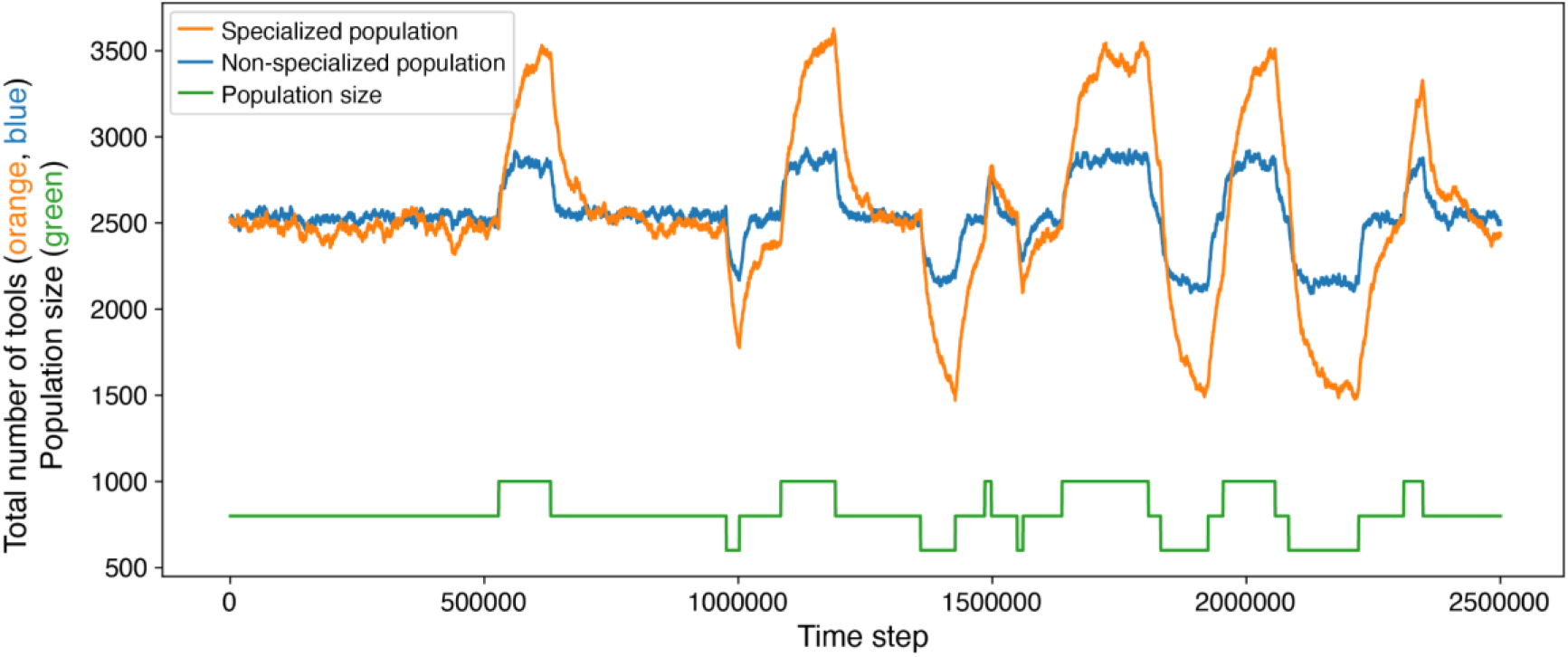
Cultural repertoires of specialized populations (here, a population divided into 10 guilds, in **orange**) are more sensitive to changes in population size than non-specialized populations (**blue**). Until time step 0.5M, population size (represented by the **green** line) is kept constant at 800 individuals, and afterwards it can vary between 600 and 1000. When population size is fixed, the cultural repertoire of the populations is similar in size (∼2500 tools). However, the specialized population is far more sensitive to fluctuations in population size (varying between 1520 and 3580 tools, while the non-specialized population varies between 2140 and 2850).

We then examined the effect of environmental switches on repertoire size of both population types (see Figure 6). We allowed a non-specialized population and a 10-guild population of 1000 to reach equilibrium repertoire sizes in a given environment. Then, we let the environment switch between two states with a fixed probability per time step. While at equilibrium, repertoire size is greater in the specialized population; however, when environments start switching, the specialized population is at a disadvantage: both the average and the minimal repertoire sizes are higher in the non-specialized population.

**Figure 6.**
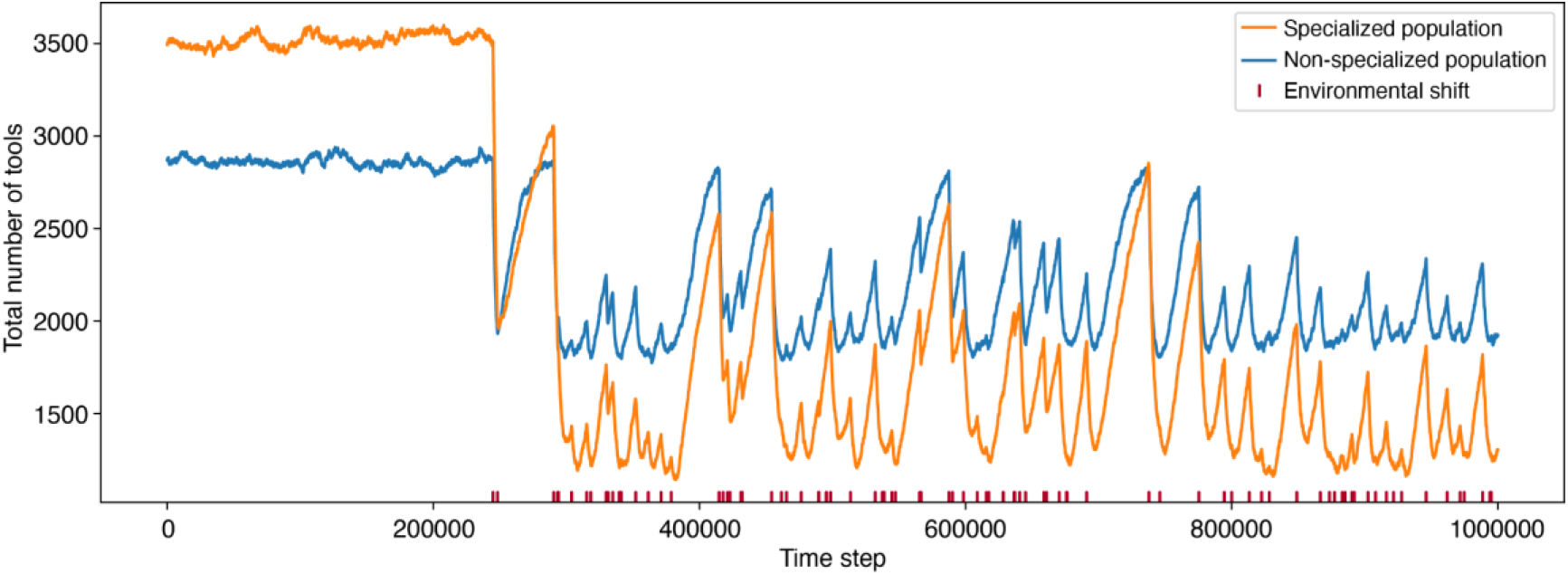
The cultural repertoire of specialized populations is more sensitive to environmental shifts than that of non-specialized populations. In a population of 1000 individuals, we find that when the environment is stable, the average cultural repertoire size of a 10-guild population (in **orange**), standing at ∼3540 tools, is higher than in the non-specialized population (in **blue**), standing at ∼2850. When environments start fluctuating (starting at time step 200000, with environmental switches marked in **dark red** on the x-axis) the average repertoire size is higher in the non-specialized population (∼2150 vs. ∼1700 tools) and the minimal repertoire is, similarly, higher in the non-specialized population (1774 vs. 1142 tools in the specialized population).

Next, we simulate the effects of a demographic bottleneck on a non-specialized population and a specialized population (Figure 7A). Our first comparison is between two populations of 1000, with a bottleneck that reduces the population size to 200 individuals. For a population of size 1000, the average repertoire size of a population divided into 10 guilds is higher (∼3540 tools) than that of a non-specialized population (∼2850 tools). After the bottleneck, on the other hand, the repertoire of the non-specialized population (∼850 tools) exceeded that of the specialized population (∼170 tools). This result indicates that the cultural loss experienced by this specialized population is bigger both in absolute and relative terms: the specialized population lost 3370 tools in comparison to 1730 in the non-specialized one, and maintained only 5% of its repertoire size, in comparison to 30%.

**Figure 7.**
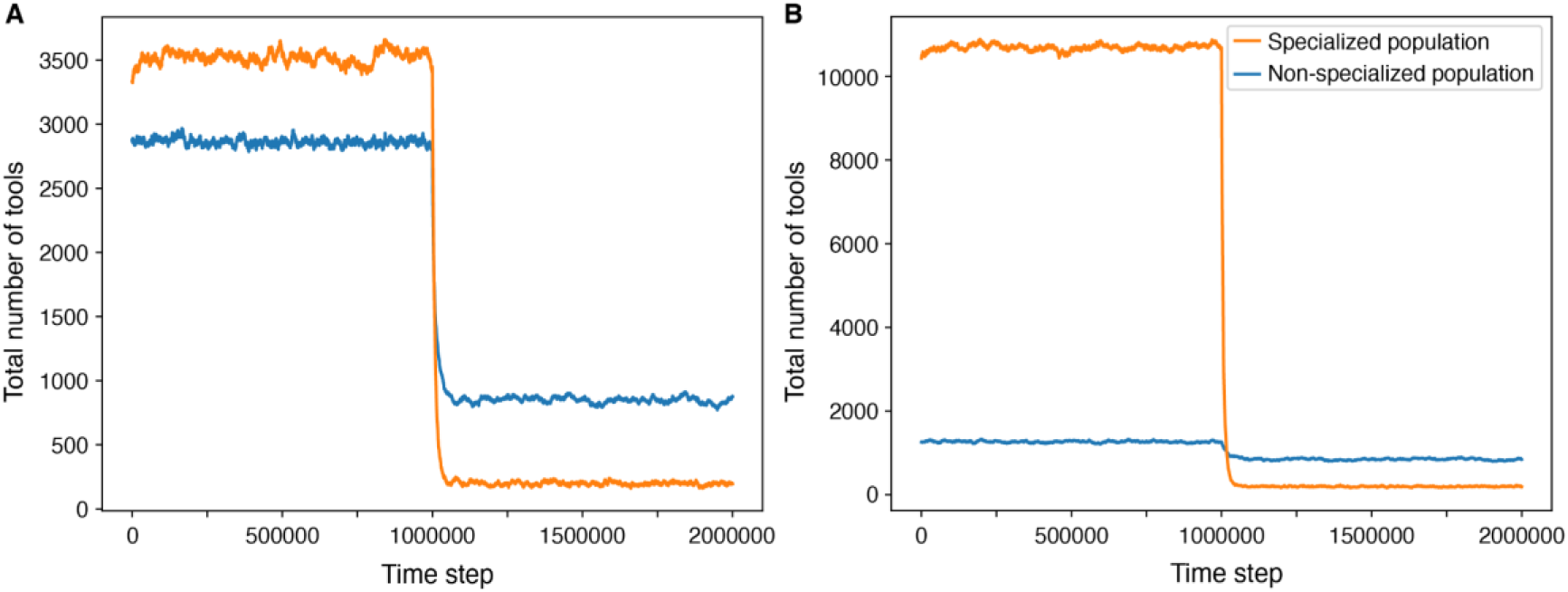
Repertoire size is reduced more drastically in specialized populations after bottlenecks. At time step 1M, the size of each population is reduced to 200 individuals, representing, for example, a migration event of a subset of the population to a new region, or a severe bottleneck following an environmental catastrophe. The **blue** line represents a non-specialized population, and the **orange** line represents a population divided into 10 guilds. Panel A simulates a reduction from 1000 individuals to 200 in both populations, in which the non-specialized population maintained ∼30% of its repertoire size while the specialized population maintained only ∼5%. Panel B simulates a reduction from 300 individuals in the non-specialized population (which maintained ∼68% of its repertoire size) and from 2500 individuals in the specialized population (which maintained ∼1%).

However, the repertoire-maximizing number of guilds for a population of 1000 is neither 1 nor 10; it is 4 (Figure 3B). In other words, under our assumptions, it may not be realistic for a population of 1000 individuals to have either 1 or 10 guilds. Thus, it may be more sensible to compare between two populations for which 1 and 10 are the optimal numbers of guilds. This means that, if populations tend to have a repertoire-maximizing number of guilds for their population size, a 10-guild population is likely to be larger than a non-specialized population. To examine whether the intuition that specialized populations tend to be larger is supported by empirical evidence, we used data on population size and the number of cultural specializations in 953 human populations (surveyed in the Ethnographic Atlas [40] and digitized in D-PLACE [41], and found a significant positive correlation (Spearman’s *ρ* = 0.53, p < 0.00001, see Figure S3). Therefore, for the specialized population, we chose a population size of 2500 (for which 10 is the repertoire-maximizing number of guilds), and for the non-specialized population a size of 300 (which is close to the highest population size for which one guild has a larger repertoire than 2 guilds). When we did this (see Figure 7B), the results became even more extreme: the specialized population maintained only 1% of its original repertoire size, while the non-specialized populations managed to save 68% of its repertoire.

## Discussion

Here, we present a model of cultural evolution that simulates the accumulation of tools in specialized and non-specialized populations under different demographic and environmental scenarios. Our model predicts that the relationship between population size and repertoire size is non-linear and can differ between non-specialized and specialized populations. For small population sizes, non-specialized populations maintain knowledge better than specialized populations and therefore reach higher average repertoire sizes. In large populations, on the other hand, specialized populations can reach higher average repertoire sizes. This is because a non-specialized population’s total repertoire size is limited by the capacity of individuals to accumulate cultural traits, while in specialized populations, each individual needs to know only a fraction of the population’s repertoire (Figure 1).

These predictions align well with prior research on human societies, which has highlighted an association between the number of specialized guilds (termed “occupational specialties” in the paper) and the overall cultural repertoire size of populations [14]. Indeed, the degree of cultural specialization in a population has been itself used as a metric of cultural complexity [14,15]. However, this previous research did not consider in detail that cultural specialization might not lead to increased cultural repertoires in smaller populations. Relatedly, an analysis of empirical research noted that a positive correlation between population size and cultural repertoire size was consistently observed in studies of food-producing populations, which tend to be larger and more sub-divided into guilds, but this correlation was not observed in studies of hunter-gatherer populations, which tend to be smaller and less specialized [13].

One question regarding cultural specialization is whether the degree of specialization is predictable for a population of a given size. In other words, what determines the number of guilds in a population? One possibility, examined in this study, is that populations will divide into the number of guilds that maximizes their cultural repertoire size. However, there could be different mechanisms that determine the observed number of guilds. In Figure S4 we explore a mechanism by which a population stabilizes on the number of guilds that can maintain a consistent repertoire size over time, which may be smaller (but are more resilient) than the maximal repertoire that the population could have reached by subdividing into more guilds. In this scenario (Figure S4), a guild that loses more than a certain percentage of its repertoire within a short time window collapses, and its members are divided between the surviving guilds. Of course, under such assumptions different numbers of guilds can emerge depending on the demographic scenario, the collapse threshold, and stochastic differences. This highlights the importance of considering not only a population’s mean repertoire but also its volatility.

Our results demonstrate that the repertoires of small guilds are particularly prone to stochastic changes. Accordingly, we suggest that populations divided into many small guilds may be less sustainable in the long term, even though their mean number of tools could be greater than that of a non-specialized population. This is true even under stable conditions, because a tool’s loss in our model—and in reality—may occur stochastically. If, in addition, environmental conditions fluctuate, rendering some tools irrelevant in certain periods of time, during which they are then more likely to be lost, this phenomenon is exacerbated: a population with small specialized guilds in a fluctuating environment may lose a disproportionate amount of its necessary cultural knowledge. A similar finding emerges with respect to a population’s ability to withstand fluctuations in its size: specialized populations with small guilds suffer disproportionate losses if population sizes fluctuate, compared to non-specialized populations.

Finally, we considered population-level bottlenecks in our simulations: extreme fluctuations in population size that may occur following natural disasters such as epidemics or extreme climatic events, as well as scenarios such as a population’s expansion to new regions. We find that bottlenecks may lead to loss of the vast majority of a population’s cultural knowledge. In contrast, the relative proportion of cultural knowledge that a non-specialized population experiences is much smaller, likely allowing such populations to maintain their subsistence patterns, social structure, and lifestyle even under a regime of repeated bottlenecks. This observation has interesting implications: it suggests, for example, that specialized populations may be less successful than non-specialized populations in spreading to new regions. Similarly, it implies that habitats that can support only a small population might not be colonizable by specialized populations.

The peopling of Australia represents a scenario in which specialization could have potentially prevented colonization. Australia was first colonized by hunter-gatherers, possibly as long as 65,000 years ago [47–49]. Since then, although some evidence exists for interactions with other populations [50,51], it seems that none of them succeeded in establishing there until the 18th century AD. Different environmental explanations have been proposed to explain what made migration to Australia possible in a specific time frame [52,53]. We propose an additional factor that may have played a role: our model suggests that while early hunter-gatherers may have been able to migrate to Australia through extreme bottlenecks without losing a substantial part of their cultural repertoire, if later societies that attempted to settle in Australia were more specialized, they would have needed to migrate in much larger numbers to be able to sustain their culture, which was not feasible until the development of advanced sailing techniques in recent millennia.

Another historical case where our model may offer a new interpretation is the collapse of the Norse settlement in Greenland. Unlike the specialized populations that might have unsuccessfully attempted to settle in Australia, farmers colonized Greenland in 986 AD and survived there successfully until disappearing mysteriously near the end of the 15th century. Many explanations have been offered to explain why this Norse settlement ceased to exist [54–56]. Some researchers (e.g. [54]) have suggested that one of the contributing factors to their collapse could have been a poor realization of the cultural repertoire of their population of origin, perhaps a result of cultural deterioration or loss of cultural elements. These researchers have also noted a decrease in the connectivity between Greenlanders and other Norse populations towards the end of their existence. The results of our model suggest that the settlement in Greenland, which was estimated to number several thousands of individuals at its highest, might not have been large enough to sustain an independent specialized population once their connection to other populations was lost. The Inuits, on the other hand, who had a culture with less occupational specialization [41], could survive even in small populations.

Several factors that may render specialized cultures unstable are not considered in our model, suggesting that its results may be conservative regarding the realistic costs of specialization. First, we used equal sized guilds. In reality, some guilds may be much smaller than others and thus particularly prone to cultural loss, and yet crucial for a population’s survival, such as traditional healers. Second, we assumed all guilds are equally represented in the bottlenecked population. In reality, after a bottleneck, populations might happen to include only few representatives from some guilds, or even none at all. Finally, subdivision of knowledge may have direct negative consequences, independent of the risk of cultural loss. For example, if knowledge of certain tools is siloed within a guild, individuals might not be aware of potentially useful tools that already exist in the population. Furthermore, a combination of two existing tools from different guilds may be useful yet remain undiscovered due to restricted knowledge sharing between them. The extent of overlap of cultural knowledge among subgroups in different societies is a promising avenue for future empirical exploration. In addition, it would be interesting to assess the degree to which tools themselves become specialized in populations divided into specialized guilds [57]. To this end, interdisciplinary studies at the intersection of theory and empirical research on this topic are particularly important [58–61].

## Supporting information

Zip of code files

## Acknowledgments

We thank Yael Sapir Ben-Oren, Sarah Saxton Strassberg, Ehud Lamm, and members of the Creanza Lab for insightful comments and discussions. OK and YB are supported by the US–Israel Binational Science Foundation (BSF) and the Israel Science Foundation (ISF; grant 1826/20). NC is supported by the National Science Foundation (NSF BCS-1918824) and the John Templeton Foundation (62187).

## Supplementary materials

**Figure S1.**
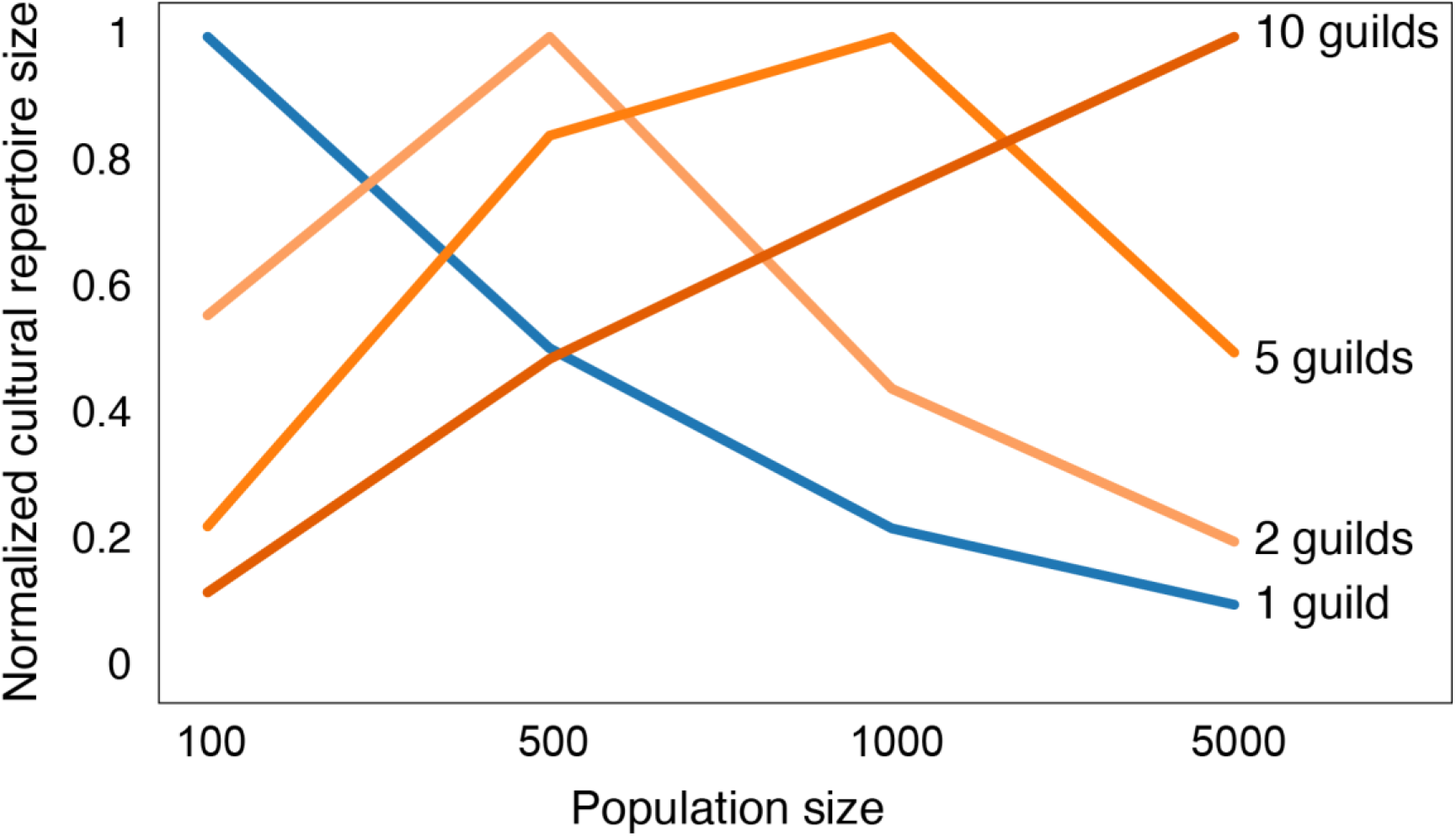
A replication of Figure 3, with a different mechanism for tool loss. Here, we examined a model in which the probability of loss of tools in a population is linearly dependent on the population size, but is also limited by a certain number, in this case 2,500 tools per guild, which is derived from the individual cognitive capacity (assuming that within a guild most individuals should know most tools). Notably, this approach yields qualitatively similar results to a model in which the population’s average cultural repertoire is an emerging property and the probability of tool loss is exponentially dependent on the existing repertoire size (Figure 3).

**Figure S2.**
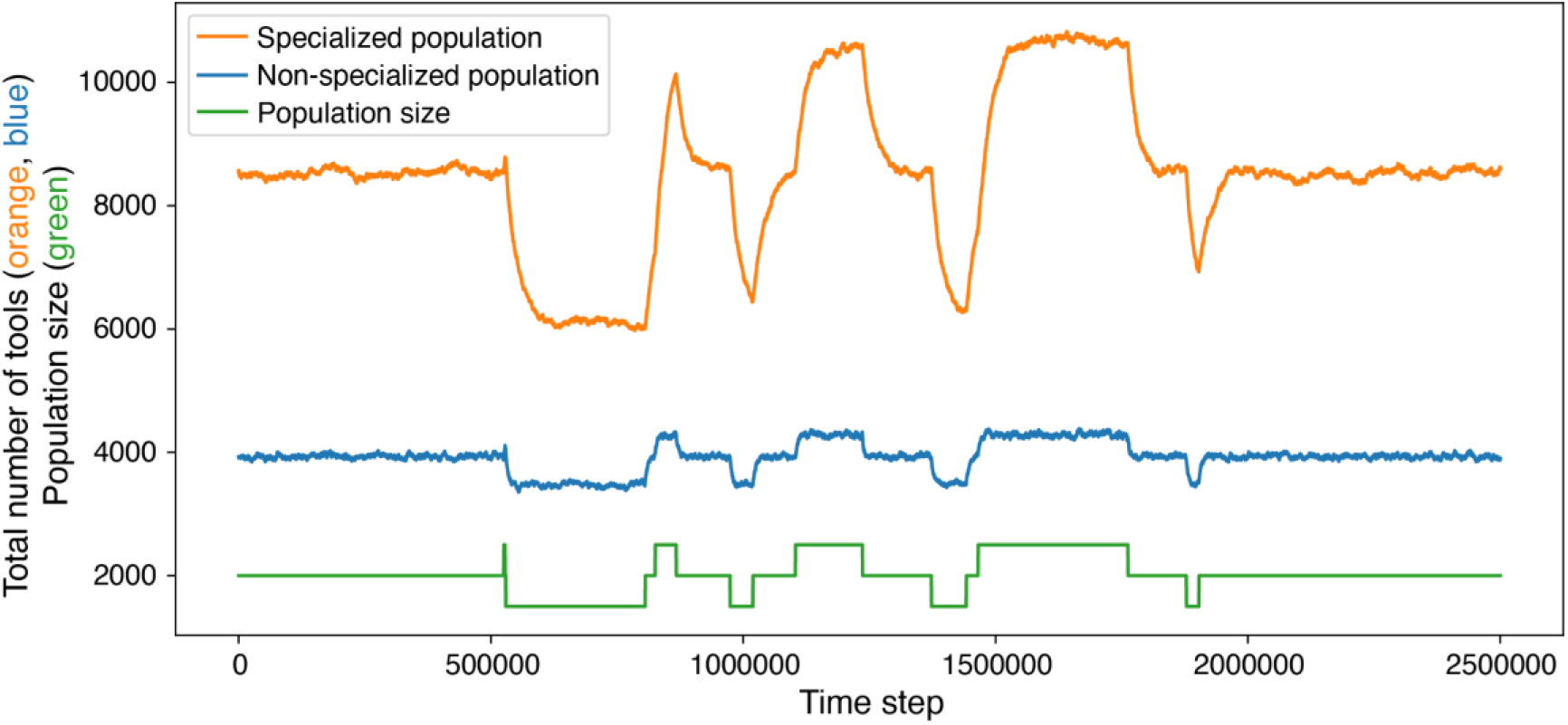
A replication of Figure 6 with a larger population size. We demonstrate here that regardless of population size, specialized populations will be more sensitive to perturbations. Here, both populations were kept at a constant size of 2000 for 0.5 million time steps, and then fluctuated between 1500 and 2500. The reduction from 2000 to 1500 resulted in a loss of 42% of the tools in the specialized population (reducing from 10750 to 8600) and only 13% in the non-specialized population (reducing from 3940 to 3480).

**Figure S3.**
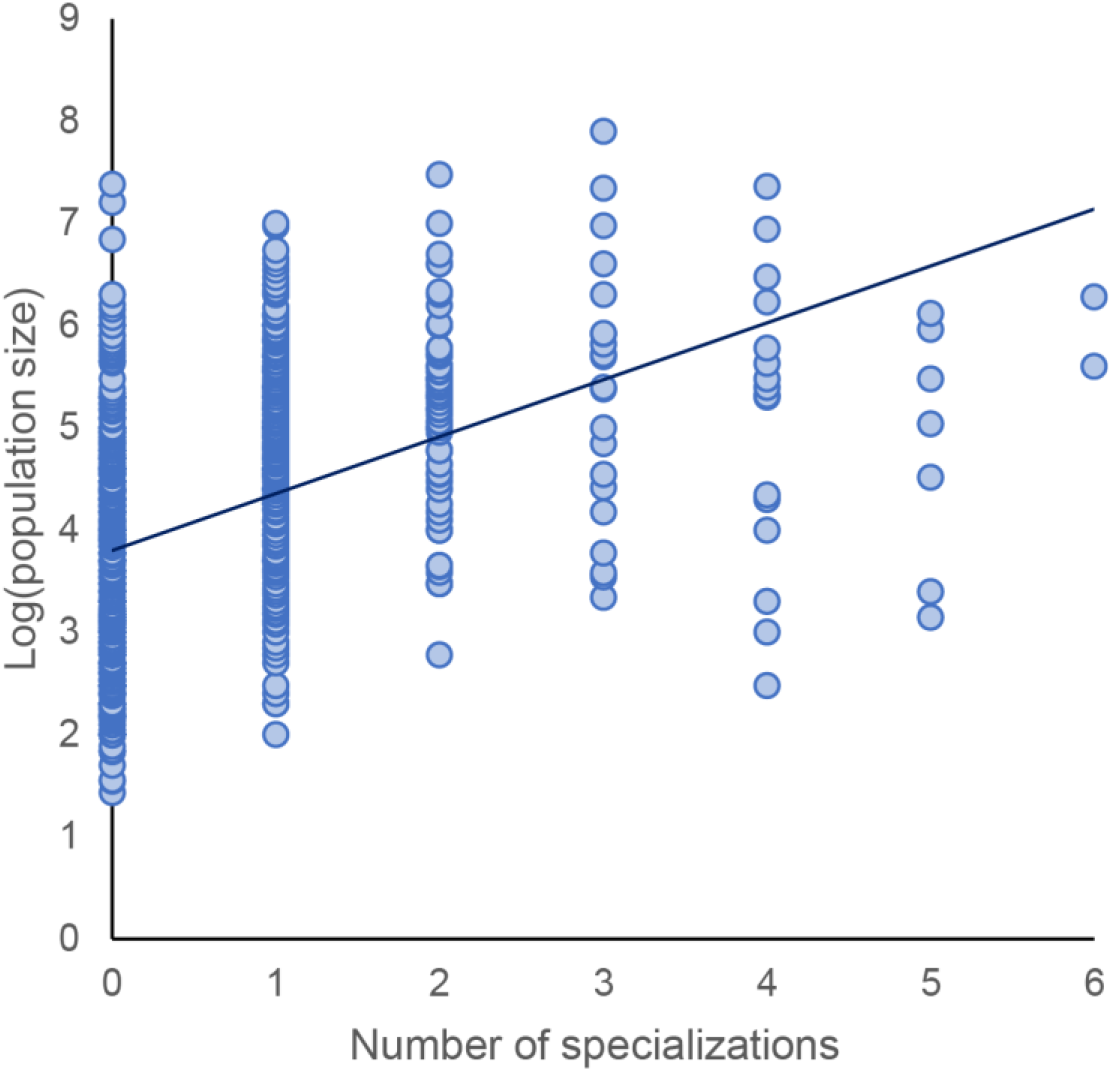
A positive correlation between population size and the number of cultural specializations across human populations worldwide. The analysis is based on data from the Ethnographic Atlas [40], digitized in D-PLACE [41]. Population size estimates were available for 953 out of the 1291 populations in the Ethnographic Atlas. For each population, we tallied how many of the eight cultural specializations listed in the database (metal working, leather working, house construction, pottery making, boat building, animal husbandry, hunting and fishing) exist as a specialized craft, indicating that related tools are known to only a subset of the population. (This is coded in the database as “Craft specialization, i.e., the activity is largely performed by a small minority of adult males or females who possess specialized skills.”) We found a significant positive correlation between these two variables (Spearman’s *ρ* = 0.53, p < 0.00001).

**Figure S4.**
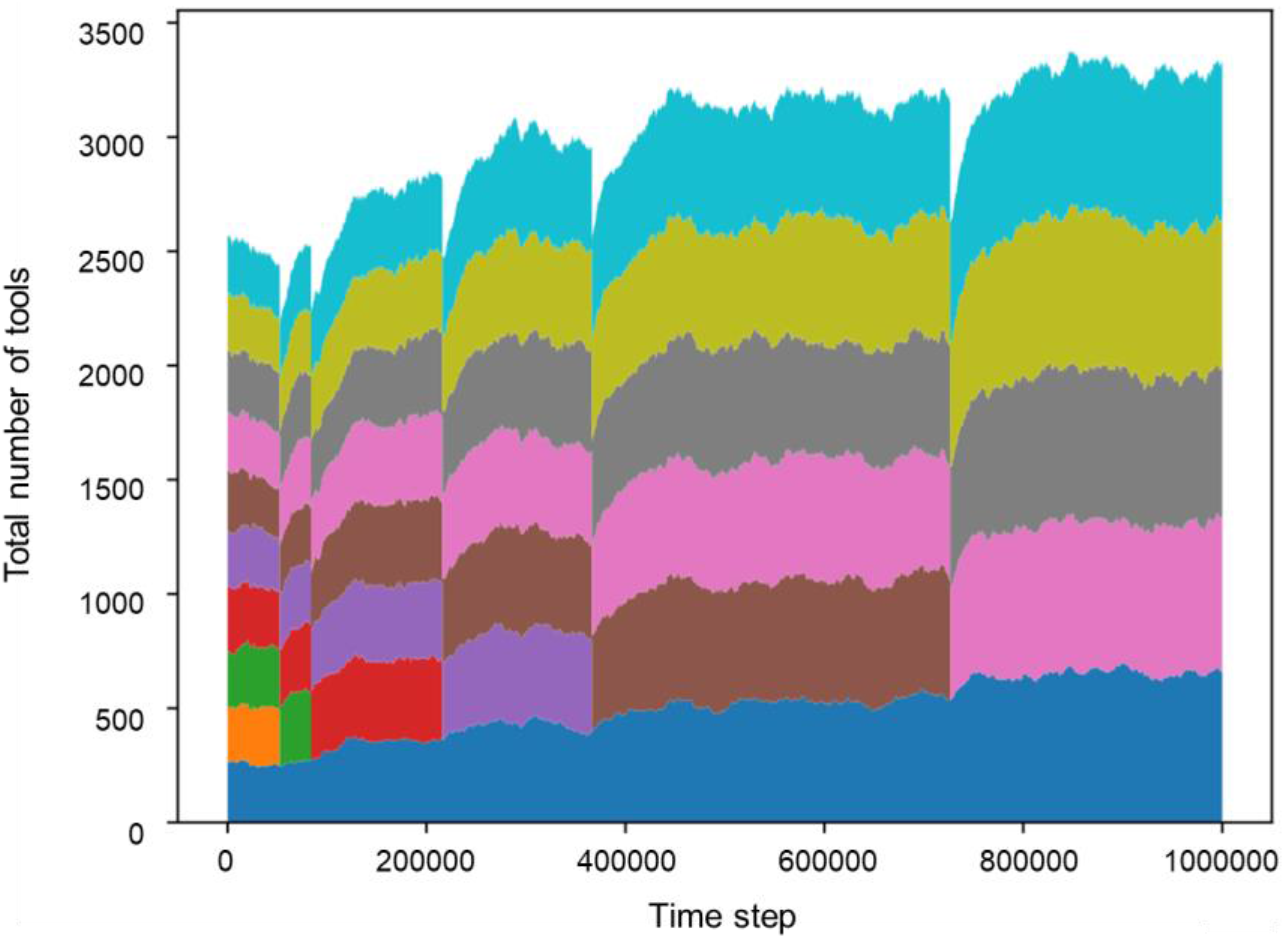
Cultural collapse due to stochasticity in a population of 800 individuals divided into 10 guilds. Each color represents the cultural repertoire of a guild. Here we defined that a guild that loses more than 10% of its repertoire size within 10,000 time steps will collapse and its members will be divided evenly between the surviving guilds. In this simulation, five of the guilds collapsed within 1 million time steps (represented in orange, green, red, purple and brown).

