## Supplementary material for "Cultural specialization as a double-edged sword: division into specialized guilds might promote cultural complexity at the cost of higher susceptibility to cultural loss": Zip of code files: Figure 6 code.py

```
#Cultural specialization model

#CCC = cultural carrying capacity

import numpy as np
import random
import matplotlib.pyplot as plt

class guild:    #we create a class called guild which has 2 attributes -
                #1. size (number of individuals)
                #2. tools (number of tools)
    def __init__(self, size, tools_a = 0, tools_b = 0, tools_both = 0):
        self.size = size
        self.tools_a = tools_a
        self.tools_b = tools_b
        self.tools_both = tools_both
        self.tools = self.tools_a + self.tools_b + self.tools_both
 
    def __repr__(self):
        return ("N = "+str(self.size)+": tools = "+str(self.tools))
    
    def tick(self,p_lucky,beta,env,new_size = False): #move to the next time unit
        
        #p_lucky = the probability of a new tool to be invented
        #an invented tool has 90% to be environment specific and 10% to be adaptive in
        #both environments.
        #beta = the beta of the binomial distribution from whice the selective value of
        #a new tool is drawn
        #loss func = the function from which the probability of tool loss is calculated
        
        if new_size != False: #if population size needs to be updated, update it
            self.size = new_size

        discovered_tools = np.random.binomial(self.size,p_lucky)
        discovered_specific = np.random.binomial(discovered_tools,0.9)
        
        if env == "a":
            lost_a = np.random.binomial(self.tools_a,(1.001**self.tools)/(500*self.size))   #for
                                                            #every tool, draw whether it was lost
            discovered_a = discovered_specific
            tools_a_s = np.random.exponential(0.1,discovered_a)   #give each discovered tool
                                                                #a selection value
            fixated_a = 0
            for s in tools_a_s: #draw which tools will be fixated
                if random.random() > 1-s:
                    fixated_a += 1
            self.tools_a = self.tools_a - lost_a + fixated_a #update the number of tools

            lost_b = np.random.binomial(self.tools_b,(1.001**self.tools)/(50*self.size))
            self.tools_b -= lost_b
            
        elif env == "b":
            lost_b = np.random.binomial(self.tools_b,(1.001**self.tools)/(500*self.size))   #for
                                                            #every tool, draw whether it was lost
            discovered_b = discovered_specific
            tools_b_s = np.random.exponential(0.1,discovered_b)   #give each discovered tool
                                                                #a selection value
            fixated_b = 0
            for s in tools_b_s: #draw which tools will be fixated
                if random.random() > 1-s:
                    fixated_b += 1
            self.tools_b = self.tools_b - lost_b + fixated_b #update the number of tools

            lost_a = np.random.binomial(self.tools_a,(1.001**self.tools)/(50*self.size))
            self.tools_a -= lost_a
            
        lost_both = np.random.binomial(self.tools_both,(1.001**self.tools)/(500*self.size))   #for
                                                            #every tool, draw whether it was lost
        discovered_both = discovered_tools - discovered_specific    #draw the number of 
                                                                    #discovered tools
        tools_both_s = np.random.exponential(0.1,discovered_both)   #give each discovered tool
                                                                #a selection value
        fixated_both = 0
        for s in tools_both_s: #draw which tools will be fixated
            if random.random() > 1-s:
                fixated_both += 1
        self.tools_both = self.tools_both - lost_both + fixated_both #update the number of tools
        self.tools = self.tools_a + self.tools_b + self.tools_both

def changing_envs(pop_size = 1000, p_lucky = 0.001,
              beta = 0.1, t_max = 1200000):

    xcoords = [] #environmental shifts
    envs = ["a"]*400000   #for 70,0000 time units, environment size will stay "a"
    
    while len(envs) < t_max: #complete the list of environments until t_max
        if random.random() < 0.00001: #with a certain probability
            if envs[-1] == "a":
                envs += ["b"]
            elif envs[-1] == "b":
                envs += ["a"]
            print("at time", len(envs)+1,"environment shifted to", envs[-1])
            xcoords += [len(envs) - 200000]
        else:
            envs += [envs[-1]]

    gen_pop = guild(pop_size) #create a generalized population
    rep_gen = [gen_pop.tools] #list of generalized population repertoire sizes across time
    
    for t in range(len(envs)): #progress the generalized population in time
        gen_pop.tick(p_lucky,beta,env = envs[t]) #update size when needed
        rep_gen += [gen_pop.tools]
        
    spec_pops = [] #a list of guilds
    rep_spec = [] #a list total repertoire sizes
    for i in range(10): #create 10 guilds
        spec_pops += [guild(pop_size/10)]
    for t in range(len(envs)): #progress specialized populations in time
        rep_at_t = []
        for pop in spec_pops: #go guild by guild
            pop.tick(p_lucky,beta,env = envs[t])
            rep_at_t += [pop.tools]
        rep_spec += [sum(rep_at_t)] #sum the repertoires of all guilds

    print("generalized population minimun", min(rep_gen[200000:t_max]),"mean:",np.mean(rep_gen[400000:t_max]))
    print("specialized population minimun", min(rep_spec[200000:t_max]),"mean:",np.mean(rep_spec[400000:t_max]))

    plt.plot(rep_gen[200000:t_max], label = "Generalized population")
    plt.plot(rep_spec[200000:t_max], label = "Specialized population")
    for shift in xcoords:
        plt.axvline(x=shift)
    plt.xlabel("Time step")
    plt.ylabel("Repertoire size")
    plt.legend(loc="upper right")
    plt.show()

def changing_envs_2(pop_size = 1000, p_lucky = 0.001,
              beta = 0.1, t_max = 1200000):
    
    envs = ["a"]*400000
    
    while len(envs) < t_max: #complete the list of environments until t_max
        if random.random() < 0.001: #with a certain probability
            if envs[-1] == "a":
                envs += ["b"]
            elif envs[-1] == "b":
                envs += ["a"]
            print("at time", len(envs)+1,"environment shifted to", envs[-1])
        else:
            envs += [envs[-1]]

    gen_pop = guild(pop_size) #create a generalized population
    rep_gen_a = [gen_pop.tools_a] #list of generalized population repertoire sizes across time
    rep_gen_b = [gen_pop.tools_b]
    rep_gen_both = [gen_pop.tools_both]
    
    for t in range(len(envs)): #progress the generalized population in time
        gen_pop.tick(p_lucky,beta,env = envs[t]) #update size when needed
        rep_gen_a += [gen_pop.tools_a]
        rep_gen_b += [gen_pop.tools_b]
        rep_gen_both += [gen_pop.tools_both]
        
    spec_pops = [] #a list of guilds
    rep_spec_a = []
    rep_spec_b = []
    rep_spec_both = []
    for i in range(10): #create 10 guilds
        spec_pops += [guild(pop_size/10)]
    for t in range(len(envs)): #progress specialized populations in time
        rep_a_at_t = []
        rep_b_at_t = []
        rep_both_at_t = []
        for pop in spec_pops: #go guild by guild
            pop.tick(p_lucky,beta,env = envs[t])
            rep_a_at_t += [pop.tools_a]
            rep_b_at_t += [pop.tools_b]
            rep_both_at_t += [pop.tools_both]
        rep_spec_a += [sum(rep_a_at_t)]
        rep_spec_b += [sum(rep_b_at_t)]
        rep_spec_both += [sum(rep_both_at_t)]
        
    labels = ["Both environmets","Environment a","Environment b"]
    
    plt.stackplot(range(len(rep_gen_a[200000:t_max])),
                  [rep_gen_both[200000:t_max], rep_gen_a[200000:t_max], rep_gen_b[200000:t_max]], labels = labels)
    plt.xlabel("Time step")
    plt.ylabel("Repertoire size")
    plt.legend(loc="upper right")
    plt.show()

    plt.stackplot(range(len(rep_spec_a[200000:t_max])),
                  [rep_spec_both[200000:t_max], rep_spec_a[200000:t_max], rep_spec_b[200000:t_max]], labels = labels)
    plt.xlabel("Time step")
    plt.ylabel("Repertoire size")
    plt.legend(loc="upper right")
    plt.show()    
    
def basic(guild_size = 100,p_lucky = 0.001, beta = 0.1,t_max = 500000):
    #run a generalized population for t_max time units and return a graph
    #of its repertoire over time
    g = guild(guild_size,0)
    tools = [g.tools]
    for t in range(t_max):
        g.tick(p_lucky,beta)
        tools += [g.tools]
    print (max(tools[200000:]))
    print (min(tools[200000:]))
    plt.plot(tools)
    plt.xlabel("Time step")
    plt.ylabel("Total number of tools")
    plt.show()
        
def optimal_n_g(n_gs = [1,2,5,10], pop_sizes = [100,500,1000,5000], p_lucky = 0.001,
                beta = 0.1,t_max = 500000,burnin = 200000): #calculate the CCC
                #for multiple population sizes and numbers of guilds

    #n_gs = a list of the numbers of guilds for which CCCs will be calculated
    #pop_sizes = a list of population sizes for which CCCs will be calculated
    #t_max = maximal time unit from which repertoire size is averaged
    #burnin = minimal time unit from which repertoire size is averaged
    results = []
    for n_g in n_gs:
        cultural_k = []
        for pop_size in pop_sizes:
            g = guild(pop_size//n_g,0)  #for multiple guilds, the size of 1 guild
                                        #used, and the carrying capacity of that
                                        #guild is then multiplied by the total number
                                        #of guilds
            tools = []
            for t in range(t_max):
                g.tick(p_lucky,beta)
                if t >= burnin:
                    tools += [g.tools]
            cultural_k += [n_g*np.mean(tools)]
        results += [cultural_k]
        print (cultural_k)

    return results

def pop_waves(pop_mean = 800, pop_max = 1000, pop_min = 600, p_lucky = 0.001,
              beta = 0.1, t_max = 2700000): #calculates the repertoire of a generalized
                #and a 10-guild population over t_max time units. starting from a
                #fixed population size (pop_mean) and then moving between pop_max
                #and pop_min
    
    current_size = pop_mean
    pop_sizes = [current_size]*700000   #for 70,0000 time units, population size will
                                        #be kept constant on pop_mean
    
    while len(pop_sizes) < t_max: #complete the list of population sizes until t_max
        while current_size < pop_max: #if population size is lower than pop_max:
            if random.random() < 0.00001: #with a certain probability
                current_size += 200 #population size can grow
            pop_sizes += [current_size]
        while current_size > pop_min: #the same for size reduction
            if random.random() < 0.00001:
                current_size -= 200
            pop_sizes += [current_size]
    gen_pop = guild(pop_sizes[0]) #create a generalized population
    rep_gen = [gen_pop.tools] #list of generalized population repertoire sizes across time
    for t in range(len(pop_sizes)): #progress the generalized population in time
        gen_pop.tick(p_lucky,beta,new_size = pop_sizes[t]) #update size when needed
        rep_gen += [gen_pop.tools]
    spec_pops = [] #a list of guilds
    rep_spec = [] #a list total repertoire sizes
    for i in range(10): #create 10 guilds
        spec_pops += [guild(pop_sizes[0]/10)]
    for t in range(len(pop_sizes)): #progress specialized populations in time
        rep_at_t = []
        for pop in spec_pops: #go guild by guild
            pop.tick(p_lucky,beta,new_size = pop_sizes[t]/10)
            rep_at_t += [pop.tools]
        rep_spec += [sum(rep_at_t)] #sum the repertoires of all guilds
    plt.plot(pop_sizes[200000:t_max], label = "Population size")
    plt.plot(rep_gen[200000:t_max], label = "Tool repertoire in the generalists' population")
    plt.plot(rep_spec[200000:t_max], label = "Tool repertoire in the specialists' population")
    plt.show()

def bn(pop_k = 1000, bn = 200, p_lucky = 0.001,
              beta = 0.1,t1 = 1100000, t2 = 1000000): #plot a population bottleneck (1000 to 200)
    #this function is very similar to "pop_waves()"; see there for more details
    gen_pop = guild(pop_k)
    rep_gen = [gen_pop.tools]
    for t in range(t1): #run generalized population until the bottleneck
        gen_pop.tick(p_lucky,beta)
        rep_gen += [gen_pop.tools]
    gen_pop.size = bn #reduce population size
    for t in range(t2): #run generalized population after the bottleneck
        gen_pop.tick(p_lucky,beta)
        rep_gen += [gen_pop.tools]
    spec_pops = []
    rep_spec = []
    for i in range(10): #create 10 guilds
        spec_pops += [guild(pop_k/10)]
    for t in range(t1):
        rep_at_t = []
        for pop in spec_pops: #run specialized population until the bottleneck
            pop.tick(p_lucky,beta)
            rep_at_t += [pop.tools]
        rep_spec += [sum(rep_at_t)]
    for pop in spec_pops:
        pop.size = bn/10 #reduce guilds' sizes
    for t in range(t2): #run specialized population after the bottleneck
        rep_at_t = []
        for pop in spec_pops:
            pop.tick(p_lucky,beta)
            rep_at_t += [pop.tools]
        rep_spec += [sum(rep_at_t)]
    #plt.plot((t1-100000)*[pop_k]+t2*[bn], label = "Population size")
    plt.plot(rep_gen[100000:2100000], label = "Tool repertoire in the generalists' population")
    plt.plot(rep_spec[100000:2100000], label = "Tool repertoire in the specialists' population")
    plt.show()

def bn_from_optimal(gen_pop_k = 300, spec_pop_k = 2500, bn = 200, p_lucky = 0.001,
              beta = 0.1,t1 = 1100000, t2 = 1000000):
    #Same as "bn()", only this time initial population sizes can be different
    #between the generalized and the specialized populations
    gen_pop = guild(gen_pop_k)
    rep_gen = [gen_pop.tools]
    for t in range(t1):
        gen_pop.tick(p_lucky,beta)
        rep_gen += [gen_pop.tools]
    gen_pop.size = bn
    for t in range(t2):
        gen_pop.tick(p_lucky,beta)
        rep_gen += [gen_pop.tools]
    spec_pops = []
    rep_spec = []
    for i in range(10):
        spec_pops += [guild(spec_pop_k/10)]
    for t in range(t1):
        rep_at_t = []
        for pop in spec_pops:
            pop.tick(p_lucky,beta)
            rep_at_t += [pop.tools]
        rep_spec += [sum(rep_at_t)]
    for pop in spec_pops:
        pop.size = bn/10
    for t in range(t2):
        rep_at_t = []
        for pop in spec_pops:
            pop.tick(p_lucky,beta)
            rep_at_t += [pop.tools]
        rep_spec += [sum(rep_at_t)]
    #plt.plot((t1-100000)*[pop_k]+t2*[bn], label = "Population size")
    plt.plot(rep_gen[100000:2100000], label = "Tool repertoire in the generalists' population")
    plt.plot(rep_spec[100000:2100000], label = "Tool repertoire in the specialists' population")
    plt.show()

def guild_collapse(n_g = 10, pop_size = 800, memory = 10000,threshold = 0.9, p_lucky = 0.001,
              beta = 0.1,t_max = 1200000, burnin = 200000):
    #run a population for "t_max" time units, where if a guild loses more than 1-"threshold"
    #of it's repertoire size within "memory" time units, it collapses and its members are
    #divided between the surviving guilds
    guilds = []
    tools = []
    surviving_g = n_g
    for g in range(n_g): #create guilds
        guilds += [guild(pop_size//n_g)]
        tools += [[guilds[g].tools]]
    for t in range(t_max):
        for i in range(n_g):
            if t > burnin: #after "burnin" timesteps, repertoires are recorded
                tools[i] += [guilds[i].tools]
            if guilds[i].size >0: #(after a guild collapses it doesn't progress)
                guilds[i].tick(p_lucky,beta)
                if t>burnin+50000 and guilds[i].tools < max(tools[i][-memory:])*threshold:
                    #after burnin+50000 guilds are allowed to collapse
                    print ("t = ", t, ". guild", i+1, "collapsed")
                    guilds[i].size = 0
                    guilds[i].tools = 0
                    surviving_g -=1
                    g_size = pop_size // surviving_g
                    for g in guilds:
                        if g.size > 0:
                            g.size = g_size
    plt.stackplot(range(len(tools[0])),tools)
    plt.show()

def choose(n_g, n): #assign n individuals to n_g guilds with equal probabilities
                    #results not expicitly discussed in the current paper
    result = [0]*n_g
    for i in range(n):
        result[random.randint(0,n_g-1)] += 1
    return result

def n_to_ccc(pop_sizes_1 = [1,5, 10, 15, 20, 25, 30, 35, 40, 45, 50],
             pop_sizes_2 = [100, 250, 400, 500, 1000, 1500, 2000, 2500, 3000, 4000, 5000, 6000, 8000, 10000], p_lucky = 0.001,
                beta = 0.1,t_max = 500000,burnin = 200000): #calculate the CCC
                #for multiple population sizes and numbers of guilds

    #n_gs = a list of the numbers of guilds for which CCCs will be calculated
    #pop_sizes = a list of population sizes for which CCCs will be calculated
    #t_max = maximal time unit from which repertoire size is averaged
    #burnin = minimal time unit from which repertoire size is average
    cccs = []
    for pop_size in pop_sizes_1:
        g = guild(pop_size,0)
        tools = []
        for t in range(t_max):
            g.tick(p_lucky,beta)
            if t >= burnin:
                tools += [g.tools]
        cccs += [np.mean(tools)]
    print (cccs)
    plt.plot(pop_sizes_1,cccs)
    plt.xlabel("Population size")
    plt.ylabel("Cultural carrying capacity")
    plt.show()
    for pop_size in pop_sizes_2:
        g = guild(pop_size,0)
        tools = []
        for t in range(t_max):
            g.tick(p_lucky,beta)
            if t >= burnin:
                tools += [g.tools]
        cccs += [np.mean(tools)]
    print (cccs)
    plt.plot(pop_sizes_1+pop_sizes_2,cccs)
    plt.xlabel("Population size")
    plt.ylabel("Cultural carrying capacity")
    plt.show()
    return cccs

def gen_vs_spec_cccs(n_gs = [1,10], pop_sizes = [10, 15, 20, 25, 30, 35, 40, 45, 50, 100, 200, 300, 400, 500, 600, 800, 1000, 1200], p_lucky = 0.001,
                beta = 0.1,t_max = 500000,burnin = 200000): #calculate the CCC
                #for multiple population sizes and numbers of guilds

    #n_gs = a list of the numbers of guilds for which CCCs will be calculated
    #pop_sizes = a list of population sizes for which CCCs will be calculated
    #t_max = maximal time unit from which repertoire size is averaged
    #burnin = minimal time unit from which repertoire size is average
    cccs = []
    for n_g in n_gs:
        g_cccs = []
        for pop_size in pop_sizes:
            g = guild(pop_size//n_g,0)
            tools = []
            for t in range(t_max):
                g.tick(p_lucky,beta)
                if t >= burnin:
                    tools += [g.tools]
            g_cccs += [np.mean(tools)*n_g]
        cccs += [g_cccs]
    print (cccs)
    plt.plot(pop_sizes,cccs[0],label = "Generalized population")
    plt.plot(pop_sizes,cccs[1],label = "Specialized population")
    plt.legend(loc="upper left")
    plt.xlabel("Population size")
    plt.ylabel("Cultural carrying capacity")
    plt.show()

    return cccs
```
