## Supplementary material for "Cultural specialization as a double-edged sword: division into specialized guilds might promote cultural complexity at the cost of higher susceptibility to cultural loss": Zip of code files: Figure S1 code.py

```
#
import numpy as np
import random
import matplotlib.pyplot as plt

class guild:
    def __init__(self, size, tools = 0):
        self.size = size
        self.tools = tools

    def __repr__(self):
        return ("N = "+str(self.size)+": tools = "+str(self.tools))
    
    def tick(self,p_lucky,beta,new_size = False): #move to the next time unit
        if new_size != False:
            self.size = new_size
        lost_tools = np.random.binomial(self.tools,0.001/self.size)
        discovered_tools = np.random.binomial(self.size,p_lucky)
        tools_s = np.random.exponential(0.1,discovered_tools)
        fixated_tools = 0
        for s in tools_s:
            if random.random() > 1-s:
                fixated_tools += 1
        self.tools = min(self.tools - lost_tools + fixated_tools,2500)

def basic(guild_size = 100,p_lucky = 0.001, p_spontloss = 0.00001, beta = 0.1,t_max = 50000):
    g = guild(guild_size,0)
    tools = [g.tools]
    for t in range(t_max):
        g.tick(p_lucky,beta)
        tools += [g.tools]
    #print (tools)
    plt.plot(tools)
    plt.xlabel("Time step")
    plt.ylabel("Total number of tools")
    plt.show()
        
def basic_w_bn(guild_size = 100, bn_size = 50, p_lucky = 0.01,
                          p_spontloss = 0.001, beta = 0.1,t1 = 50000,t2 = 50000, t3 = 50000):
    g = guild(guild_size,0)
    tools = [g.tools]
    for t in range(t1):
        g.tick(p_lucky,beta)
        tools += [g.tools]
    g.size = bn_size
    for t in range(t2):
        g.tick(p_lucky,p_spontloss,beta)
        tools += [g.tools]
    g.size = guild_size
    for t in range(t1):
        g.tick(p_lucky,p_spontloss,beta)
        tools += [g.tools]
    #print (tools)
    plt.plot(tools)
    plt.xlabel("Time step")
    plt.ylabel("Total number of tools")
    plt.show()

def five_guilds(n_guilds = 5, guild_size = 20, p_lucky = 0.01, p_spontloss = 0.00001, beta = 0.1,t_max = 50000):
    guilds = []
    tools = []
    current_tools = []
    for i in range(n_guilds):
        new_guild = guild(guild_size,0)
        guilds += [new_guild]
        current_tools += [new_guild.tools]
    tools += [current_tools]
    for t in range(t_max):
        current_tools = []
        for g in guilds:
            g.tick(p_lucky,p_spontloss,beta)
            current_tools += [g.tools]
        tools += [current_tools]
    matrix = np.array(tools)
    g1 = matrix[:,0]
    g2 = matrix[:,1]
    g3 = matrix[:,2]
    g4 = matrix[:,3]
    g5 = matrix[:,4]

    #print (tools)
    plt.stackplot(range(len(tools)),g1,g2,g3,g4,g5)
    plt.xlabel("Time step")
    plt.ylabel("Total number of tools")
    plt.show()
    return None

def five_guilds_w_bn(n_guilds = 5, guild_size = 20, bn_size = 10, p_lucky = 0.01,
                     p_spontloss = 0.001, beta = 0.1,t1 = 50000,t2 = 50000, t3 = 50000):
    guilds = []
    tools = []
    current_tools = []
    for i in range(n_guilds):
        new_guild = guild(guild_size,0)
        guilds += [new_guild]
        current_tools += [new_guild.tools]
    tools += [current_tools]
    for t in range(t1):
        current_tools = []
        for g in guilds:
            g.tick(p_lucky,p_spontloss,beta)
            current_tools += [g.tools]
        tools += [current_tools]
        
    for g in guilds:
        g.size = bn_size

    for t in range(t2):
        current_tools = []
        for g in guilds:
            g.tick(p_lucky,p_spontloss,beta)
            current_tools += [g.tools]
        tools += [current_tools]

    for g in guilds:
        g.size = guild_size

    for t in range(t3):
        current_tools = []
        for g in guilds:
            g.tick(p_lucky,p_spontloss,beta)
            current_tools += [g.tools]
        tools += [current_tools]
        
    matrix = np.array(tools)
    g1 = matrix[:,0]
    g2 = matrix[:,1]
    g3 = matrix[:,2]
    g4 = matrix[:,3]
    g5 = matrix[:,4]

    #print (tools)
    plt.stackplot(range(len(tools)),g1,g2,g3,g4,g5)
    plt.xlabel("Time step")
    plt.ylabel("Total number of tools")
    plt.show()
    return None

def optimal_n_g(n_gs = [1,2,5,10], pop_sizes = [10,50,100,200,300,400,500], p_lucky = 0.001,
                beta = 0.1,t_max = 300000,burnin = 100000):
    results = []
    for n_g in n_gs:
        cultural_k = []
        for pop_size in pop_sizes:
            g = guild(pop_size//n_g,0)
            tools = []
            for t in range(t_max):
                g.tick(p_lucky,beta)
                if t >= burnin:
                    tools += [g.tools]
            cultural_k += [n_g*np.mean(tools)]
        results += [cultural_k]

    matrix = np.array(results)
    print (matrix)
    curve1 = matrix[0]
    curve2 = matrix[1]
    curve3 = matrix[2]
    curve4 = matrix[3]

    plt.plot(pop_sizes,curve1)
    plt.plot(pop_sizes,curve2)
    plt.plot(pop_sizes,curve3)
    plt.plot(pop_sizes,curve4)
    plt.show()

    return None

def pop_waves(pop_mean = 800, pop_max = 1000, pop_min = 600, p_lucky = 0.001,
              beta = 0.1,t_max = 2100000):
    
    current_size = pop_mean
    pop_sizes = [current_size]*600000
    
    while len(pop_sizes) < t_max:
        while current_size < pop_max:
            if random.random() < 0.00001:
                current_size += 200
            pop_sizes += [current_size]
        while current_size > pop_min:
            if random.random() < 0.00001:
                current_size -= 200
            pop_sizes += [current_size]
    gen_pop = guild(pop_sizes[0])
    rep_gen = [gen_pop.tools]
    for t in range(len(pop_sizes)):
        gen_pop.tick(p_lucky,beta,new_size = pop_sizes[t])
        rep_gen += [gen_pop.tools]
    spec_pops = []
    rep_spec = []
    for i in range(10):
        spec_pops += [guild(pop_sizes[0]/10)]
    for t in range(len(pop_sizes)):
        rep_at_t = []
        for pop in spec_pops:
            pop.tick(p_lucky,beta,new_size = pop_sizes[t]/10)
            rep_at_t += [pop.tools]
        rep_spec += [sum(rep_at_t)]
    plt.plot(pop_sizes[100000:2100000], label = "Population size")
    plt.plot(rep_gen[100000:2100000], label = "Tool repertoire in the generalists' population")
    plt.plot(rep_spec[100000:2100000], label = "Tool repertoire in the specialists' population")
    plt.show()

def bn(pop_k = 5000, bn = 500, p_lucky = 0.001,
              beta = 0.1,t1 = 1100000, t2 = 1000000):
    
    gen_pop = guild(pop_k)
    rep_gen = [gen_pop.tools]
    for t in range(t1):
        gen_pop.tick(p_lucky,beta)
        rep_gen += [gen_pop.tools]
    gen_pop.size = bn
    for t in range(t2):
        gen_pop.tick(p_lucky,beta)
        rep_gen += [gen_pop.tools]
    spec_pops = []
    rep_spec = []
    for i in range(10):
        spec_pops += [guild(pop_k/10)]
    for t in range(t1):
        rep_at_t = []
        for pop in spec_pops:
            pop.tick(p_lucky,beta)
            rep_at_t += [pop.tools]
        rep_spec += [sum(rep_at_t)]
    for pop in spec_pops:
        pop.size = bn/10
    for t in range(t2):
        rep_at_t = []
        for pop in spec_pops:
            pop.tick(p_lucky,beta)
            rep_at_t += [pop.tools]
        rep_spec += [sum(rep_at_t)]
    #plt.plot((t1-100000)*[pop_k]+t2*[bn], label = "Population size")
    plt.plot(rep_gen[100000:2100000], label = "Tool repertoire in the generalists' population")
    plt.plot(rep_spec[100000:2100000], label = "Tool repertoire in the specialists' population")
    plt.show()

def guild_collapse(n_g = 10, pop_size = 800, memory = 10000,threshold = 0.9, p_lucky = 0.001,
              beta = 0.1,t_max = 600000, burnin = 100000):
    guilds = []
    tools = []
    surviving_g = n_g
    for g in range(n_g):
        guilds += [guild(pop_size//n_g)]
        tools += [[guilds[g].tools]]
    for t in range(t_max):
        for i in range(n_g):
            if t > burnin:
                tools[i] += [guilds[i].tools]
            if guilds[i].size >0:
                guilds[i].tick(p_lucky,beta)
                if t>burnin and guilds[i].tools < max(tools[i][-memory:])*threshold:
                    print ("t = ", t, ". guild", i+1, "collapsed")
                    guilds[i].size = 0
                    guilds[i].tools = 0
                    surviving_g -=1
                    g_size = pop_size // surviving_g
                    for g in guilds:
                        if g.size > 0:
                            g.size = g_size
    plt.stackplot(range(len(tools[0])),tools)
    plt.show()

def guild_collapse_waves(n_g = 10, pop_size = 5000, memory = 10000,threshold = 0.9, p_lucky = 0.001,
              beta = 0.1,t_max = 600000, burnin = 100000):
    guilds = []
    tools = []
    surviving_g = n_g
    pop_sizes = [pop_size]*burnin
    
    while len(pop_sizes) < t_max:
        while current_size < pop_max:
            if random.random() < 0.00001:
                current_size += 200
            pop_sizes += [current_size]
        while current_size > pop_min:
            if random.random() < 0.00001:
                current_size -= 200
            pop_sizes += [current_size]
    
    for g in range(n_g):
        guilds += [guild(pop_size//n_g)]
        tools += [[guilds[g].tools]]
    for t in range(t_max):
        for i in range(n_g):
            if t > burnin:
                tools[i] += [guilds[i].tools]
            if guilds[i].size >0:
                guilds[i].tick(p_lucky,beta)
                if t>burnin and guilds[i].tools < max(tools[i][-memory:])*threshold:
                    print ("t = ", t, ". guild", i+1, "collapsed")
                    guilds[i].size = 0
                    guilds[i].tools = 0
                    surviving_g -=1
                    g_size = pop_size // surviving_g
                    for g in guilds:
                        if g.size > 0:
                            g.size = g_size
    plt.stackplot(range(len(tools[0])),tools)
    plt.show()
```
